# Inferring species interactions from ecological survey data: a mechanistic approach to predict quantitative food webs of seed-feeding by carabid beetles

**DOI:** 10.1101/2020.11.09.375402

**Authors:** Michael J.O. Pocock, Reto Schmucki, David A. Bohan

## Abstract

1. Ecological networks are valuable for ecosystem analysis but their use is often limited by a lack of data because many types of ecological interaction, e.g. predation, are short-lived and difficult to observe or detect. There are different methods for inferring the presence of interactions, which we lack methods to predict interaction strengths and so use weighted network analysis.
2. Here, we develop a trait-based approach suitable for creating quantitative networks, i.e. with varying interaction strengths. We developed the method for seed-feeding carabid ground beetles (Coleoptera: Carabidae) although the principles can be applied to other interactions.
3. We used existing literature data from experimental seed-feeding trials to predict a per-individual interaction cost index based on carabid and seed size with frequency-dependent prey selection and assuming bottom up control. This was scaled up to the population level to create predicted inferred weighted networks using the abundance of carabids and seeds in samples from arable fields and energetic intake rates of carabids from the literature. From these weighted networks, we also calculated a novel measure of predation pressure.
4. We applied it to existing ecological survey data from 255 arable fields with carabid data from pitfall traps and plant seeds from seed rain traps. Analysis of these inferred networks led to testable hypotheses about how networks and predation pressure varied amongst fields.
5. Inferred networks are valuable because (i) they provide null models for the structuring of food webs to test against empirical species interaction data, e.g. DNA analysis of carabid gut regurgitates, and (ii) they allow weighted inferred networks to be constructed whenever we can estimate interactions between species and have ecological census data available. This would permit network analysis even at times and in places when interactions were not directly assessed.

## Introduction

Networks are a valuable tool for understanding the structure and dynamics of ecosystems (Kaiser-Bunbury & Blüthgen, 2015; Ma et al., 2019; Pocock, Evans, & Memmott, 2012; Tylianakis, Laliberté, Nielsen, & Bascompte, 2010). Ecological networks are constructed from information on biotic interactions, e.g. species-species interactions, and provide a whole system approach to understanding changes in biodiversity, the resilience of biodiversity to environmental change and the provision of ecosystem function (Heleno et al., 2014). Weighted networks are especially valuable in ecological networks because many ecosystem functions are influenced by interaction frequency or importance (Kaiser-Bunbury & Blüthgen, 2015). Despite the value of a network approach to studying ecological systems, we are often severely limited by the lack of empirical data on biotic interactions: this has been termed the ‘Eltonian shortfall’ (Hortal et al., 2015). In the past researchers have sought to address this shortfall by using different methods to infer the presence of interactions from data on species presence, but we need methods to infer quantitative interactions and so gain the benefits from using weighted network analysis.

The lack of data on species interactions is because many interactions are relatively brief (e.g. an animal consuming an arthropod) or cryptic (e.g. predation occurring at night or underground). This means that interactions are more difficult to sample than species. As a consequence, many ecological networks studies are based on (i) interactions that are comparatively easy to record, e.g. direct observations of pollinating insects visiting flowers (Memmott, 1999) and analysis of gut contents of ‘gulp predators’ such as fish (Gray et al., 2014); or (ii) interactions that are long-lasting, e.g. parasitoids and their hosts (Müller, Adriaanse, Belshaw, & Godfray, 1999) and phytophagous insects and their food plants, both of which can be identified using rearing or DNA-based methods (Evans et al. 2016). Even in these examples, adequate sampling the network of interactions is far more costly (in time and/or resources) than sampling the organisms alone (Banašek-Richtera, Cattina, & Bersier, 2004). This means that many types of biotic interaction are under-represented in ecological research, despite being of interest and high functional importance.

There are many datasets where assemblages of potentially-interacting species (e.g. predators and prey) have been sampled using standard ecological census techniques, but since the interactions themselves have not been sampled they cannot be used for network analyses. If it was possible to infer these interactions, these datasets would provide a vast resource about changes in (inferred) networks over time and space (Delmas et al., 2019) and hence on ecosystems and their dynamics. The inference of biotic interactions is an active area of research and broadly four approaches have been used in the past.

1. *Using databases of previously-recorded interactions* and applying these to datasets of co-occurring species. This relatively simple approach has been used for a range of taxa (Gray et al., 2014; Muñoz, Trøjelsgaard, & Kissling, 2019; Redhead et al., 2018), and it makes the assumption that if two species were recorded to interact at one time and place then they will always interact when they co-occur. This also depends on the availability and comprehensiveness of information on recorded interactions (Poelen, Simons, & Mungall, 2014). Databases could be biased towards common interactions, because they are most frequently observed, or towards uncommon interactions, because they are most frequently reported (because the observer deems them to be noteworthy).
2. *Co-occurrence analysis* is used to statistically define associations from samples obtained at the same time and place. This is particularly useful for datasets of many discrete samples, e.g. DNA meta-barcoding of species from environmental samples (Lima-Mendez et al., 2015; Vacher et al., 2016) or gene regulatory networks from gene-expression microarray data (Marbach et al., 2012). It has been applied to biodiversity data, where citizen science provides a large number of co-occurence records (Milns, Beale, and Smith 2010). One challenge with this approach is that when an association does occur, further information is required to determine the type of interaction (Faust & Raes, 2012; Freilich, Wieters, Broitman, Marquet, & Navarrete, 2018), e.g. an association between two species could be due to predation, mutualism or shared resource use.
3. *Inductive machine-learning approaches* are data-intensive but only require simple rules to be provided (e.g. small species are preyed upon by larger species) and the networks are learned from data. For example, inductive machine learning has been used with time series of species abundance data from agro-ecosystems to successfully identify previously under-recorded predatory interactions, namely: spiders predating small carabid ground beetles (Bohan, Caron-Lormier, Muggleton, Raybould, & Tamaddoni-Nezhad, 2011).
4. *Trait-based approaches* have often been used when there is an allometric relationship between size of predator and prey, e.g. gape size limits prey selection (Gravel, Poisot, Albouy, Velez, & Mouillot, 2013; Petchey, Beckerman, Riede, & Warren, 2008; Sebastián-González, Pires, Donatti, Guimarães, & Dirzo, 2017; Williams, Anandanadesan, & Purves, 2010), but they can be used for a range of other interaction proxies, such as phylogenetic or niche distance (Morales-Castilla, Matias, Gravel, & Araújo, 2015).

These inference approaches have different benefits, depending on the datasets available and the ambitions of the researchers (Faisal, Dondelinger, Husmeier, & Beale, 2010). However, the inferred networks are typically limited to presence (or probability) of interactions, rather than the interaction weighted. Quantifying the weights, or strengths, of interactions is important. Firstly, for a practical sense interaction strengths are less biased to sampling effects and so weighted network analysis will provide insights into ecological function and resilience that are more robust to sampling biases than unweighted network analysis (Bersier, Banašek-Richter, & Cattin, 2002). Secondly, measures of interaction strengths provide information on the functional importance of interactions (Vazquez et al., 2007) therefore, in general, weighted interactions are likely to be more ecologically meaningful than simply the presence of an interaction.

Here we considered weed seeds in arable fields and their predation by carabids (ground beetles, Coleoptera: Carabidae). Carabids are important multi-functional predators in agro-ecosystems (Honek, Martinkova, & Jarosik, 2003; Ma et al., 2019). Many are important seed feeders with seeds either as part or all of their diet, depending on the species (Honek et al., 2003; Kulkarni, Dosdall, & Willenborg, 2015). Practically, seed-feeding carabids provide a valuable ecosystem service in helping to regulate weed seeds in arable fields (Bohan, Boursault, Brooks, & Petit, 2011; Petit et al., 2018). While carabids and seeds can be sampled relatively straightforwardly with standard ecological census techniques (Brooks et al., 2003; Heard et al., 2003), it is much harder to detect their seed-feeding interactions. Here we develop a mechanistic model of prey preference (frequency-dependent foraging) to infer weighted interactions between seeds and carabids and apply it to a large dataset of weed seed and carabid abundances.

## Methods

### Developing a mechanistic trait-based model to infer weighted food webs

Here we develop a mechanistic model of predator-prey interactions of carabid ground beetles preying upon weed seeds at the soil surface of arable fields. The fundamental variable needed to construct weighted ecological interaction networks is the frequency of each prey in the diet of each predator present in the field (*F*_*ij*_). This relies on predicting the foraging behaviour and prey selection of the predators. In this case, we needed a model of prey selection based on *relative* search and handling time (so that we could use data from multiple feeding trials in the literature). We used a model for frequency-dependent prey selection (Gendron, 1987) in which the frequency (*F*_*i*_) of a prey item *i* in the diet of a predator is the normalised function of its ‘risk index’ (*r*_*i*_) and its density (*D*_*i*_). The calculation of *F*_*ij*_ therefore requires calculation of the risk index (*r*_*ij*_), which itself requires estimation of an interaction cost index (*h*_*ij*_). Having obtained estimates of the frequency of each seed in the diet of each carabid, we scaled up the estimates to construct a whole inferred network with currently available data on seed and carabid density in arable fields (Fig. 1; Table 1). This process is described in detail below.

**Fig. 1.**
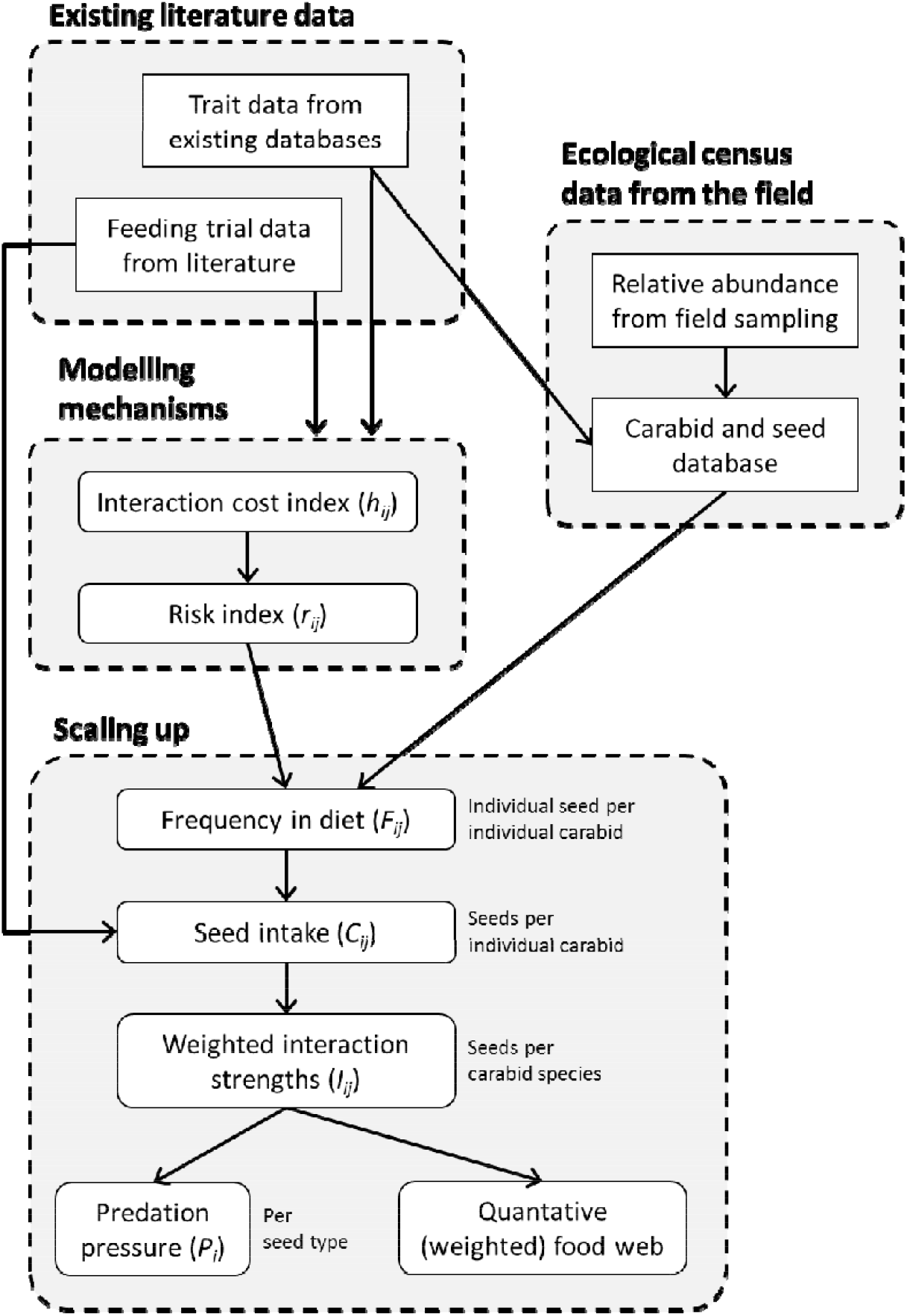
Summary of the process to use data from ecological sampling and from the literature to scale up to inferred food webs.

**Table 1.**
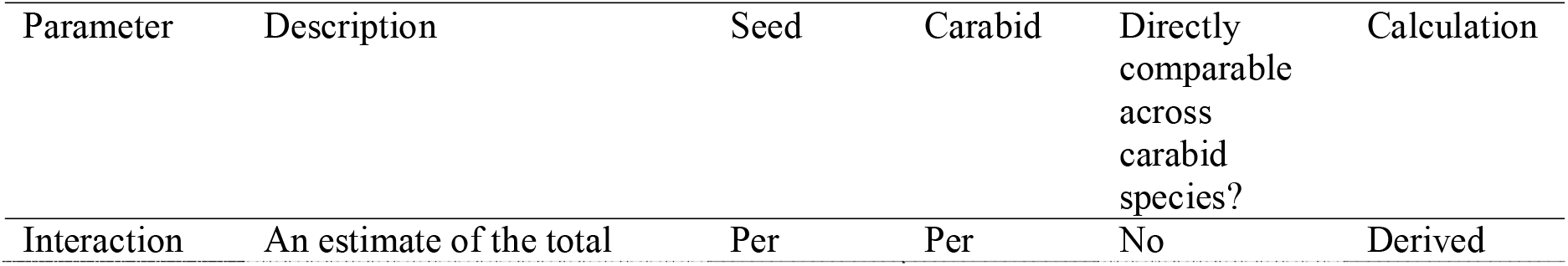
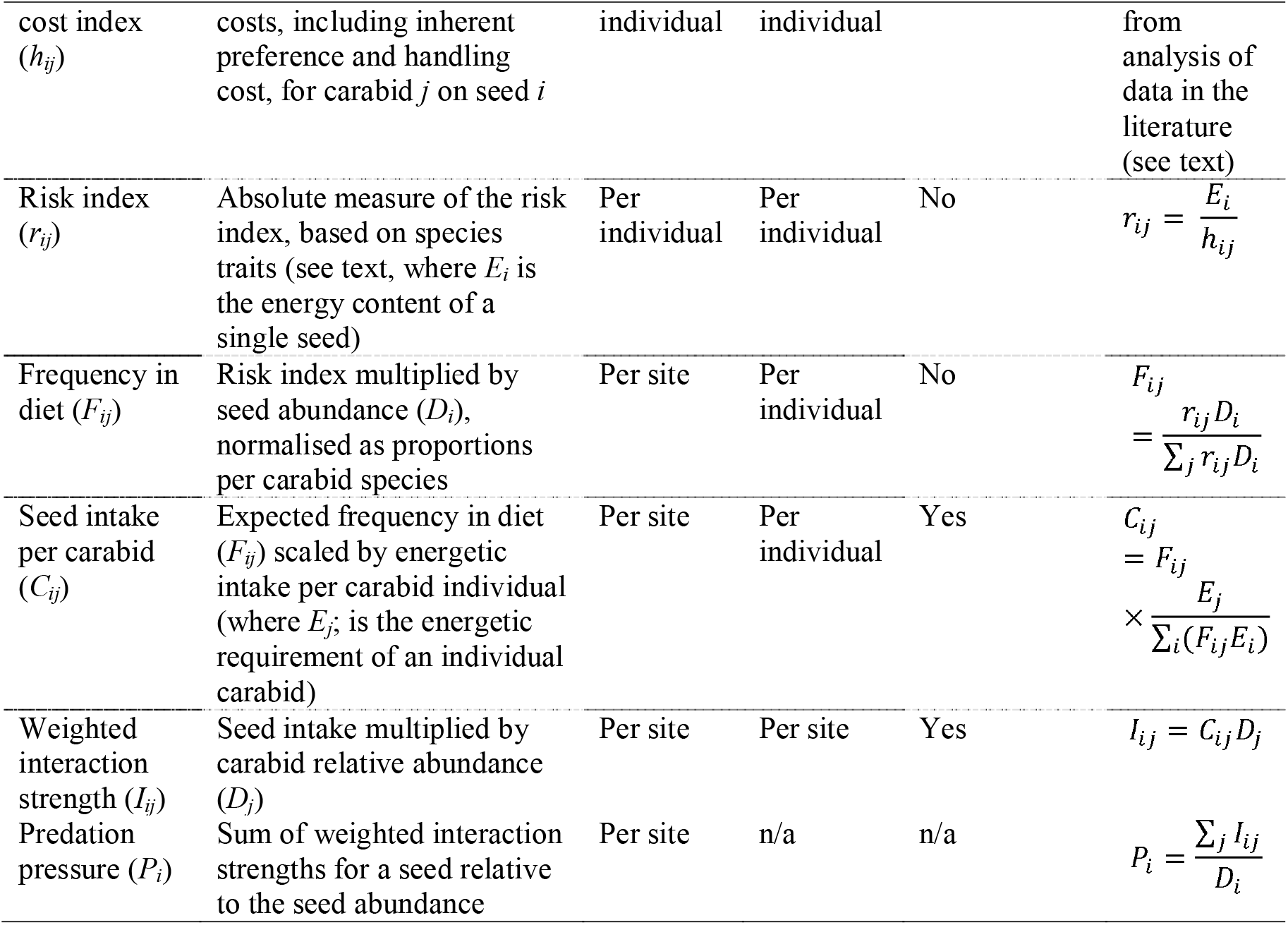
Parameters in the conversion of risk indices to interaction strengths for construction of weighted food webs of seed-feeding carabids. In our example, we consider *i* = seed genus and *j* = carabid species.

The data we used to calculate the interaction cost index (*h*) were obtained from published ‘cafeteria experiments’, which are choice experiments in which the consumption of prey by predators is tested in experimental arenas. These have been used to assess carabid prey preferences for seeds (Honek, Martinkova, Saska, & Pekar, 2007; Petit, Boursault, & Bohan, 2014) and invertebrates (Lang & Gsödl, 2008). We obtained data from cafeteria experiments for two species of carabid feeding on 64 species of seed (Honek et al., 2003) and for five species of carabid feeding on ten species of seed (Petit et al., 2014). In these studies individual beetles were provided with seeds of seven (Honek et al., 2003) or ten (Petit et al., 2014) different species of seed in experimental arenas. Researchers daily recorded the number of seeds that had been consumed. For Honek et al. (2003), the data on time to remove 50% of seeds (CT_50_) were digitised from Figure 4 in and converted to daily consumption rate (*x*_*i*_): *x*_*i*_ = (0.5 × number of seeds used in experiment) / CT_50_.

We assumed that the number of seeds consumed was proportional to the ‘risk index’ of that seed species for that carabid species (*x*_*i*_ ∝ *r*_*i*_). We combined this with the definition of the risk index: *r*_*i*_ = *E*_*i*_/*h*_*i*_, where *E*_*i*_ is energy content of the prey item *i* and *h* is its ‘handling cost’ for predator *j* (Tinbergen, 1960). The parameter *h* describes the totality of the ‘costs’ that are included in the risk index (e.g. handling time, innate preference of the predator for the prey and so on) and so, hereafter, this parameter is termed the interaction cost index. By re-arrangement, we can define: *h*_*i*_ = *E*_*i*_/*x*_*i*_., which means that *h*_*i*_ was comparable within but not between different experiments. We then used predator and prey size to model *h* because these traits are major determinants of weed seed preferences (Kulkarni, Dosdall, & Willenborg, 2015). Specifically modelled the interaction cost index (*h*_*ij*_) as a function of the interaction of seed size (cube root of seed mass) and carabid size (log-transformed body mass), and included the study as a random effect:

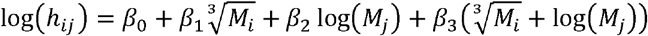

There were few very large seeds (>5mg), but their predicted interaction costs were high (Fig. S1) so we weighted the data points by the inverse of the cube-root of seed mass 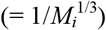 to reduce their influence on the results. To calculate the risk index for each carabid and seed combination, we used the interaction cost index and the energetic content of the seed (Table 1). By using a trait-based approach to estimate *h* from data in the literature meant that we could estimate the risk index (*r*) for carabid and seed species (based on their mass) even if they were not included in the published cafeteria experiments.

Having obtained the risk index for each carabid-seed combination, we scaled this by the density of the seeds (*D*_*i*_) and normalised it to calculate the predicted frequency (*F*_*ij*_) of seed *i* in the diet of an individual carabid of species *j*. We then needed information on the energy intake of carabids in different feeding guilds. Since the seed predators in this study were all closely related carabid species, they are likely to have the same energetic intake (per gram of body mass) of seeds, once their feeding guild was taken into account. Several sources were used to categorise British carabid species into feeding guilds (Brooks et al., 2012; Harvey, van der Putten, Turin, Wagenaar, & Bezemer, 2008; Luff, 2007): species in Tribes Harpalini and Zabrini were classed as granivorous; species in Tribe Sphodrini, Tribe Trechini, and genera *Pterostichus* and *Poecilus* (from Tribe Pterostchini) and *Agonum* (from Tribe Platynini) were classed as omnivorous; all other species were classed as obligate carnivores. Published data provided information on the daily energy intake of seeds (mg of seeds per mg carabid body mass). (Note that these are different experiments to the choice experiments used to estimate the handling costs, explained above.) In Honek et al. (2003; their Table 2), 23 species of carabid were given *Cirsium arvense* (1.3mg) and *Capsella bursa-pastoralis* (0.1mg) *ad lib*. in separate experiments; we used data for the seed with the maximum consumption rate. In Petit et al. (2014; their Table 1), five species of carabid were each given ten species of seed (0.1-8.9mg) together in a cafeteria experiment. We totalled the daily mass of seeds consumed and modelled the average feeding rate (in mg seeds per mg body mass) for species in the different guilds with a mixed-effects model with feeding guild as a fixed effect, and species and source of data as random effects.

We scaled up the literature-derived average feeding rate to the per-individual energetic requirement of carabids (i.e. *E*_*j*_ = feeding rate (mg per mg) × *M*_*j*_), and then converted this from the *mass* of seeds to the *number* of seeds of type *i* to give the seed intake per carabid (*C*_*ij*_ = number of seeds of species *I* consumed by an individual of carabid *j*). We then multiplied this by the abundance of carabids of species *j* to give the weighted interaction strength (*I*_*ij*_) and so combined these to construct a weighted, inferred food web. Finally, we calculated a predation pressure ratio (*P*_*i*_) for each seed type in each field to provide an assessment of the predicted intensity of seed predation based on the inferred food web.

### Applying the inference of interactions to ecological census data of weed seeds and carabids in arable fields

We applied this analysis pipeline to construct inferred food webs with data on seed and carabid abundance from 255 arable fields. (See http://doi.org/10.5281/zenodo.4252783 for R code and data to replicate this analysis). The data we used were obtained from the Farm Scale Evaluation (FSE) of genetically modified crops. The FSE was a split-field experiment comparing the impact of four genetically modified crops to the same crops cultivated conventionally (Firbank et al., 2003). The 255 fields (67 fields of spring oil seed rape, 65 of winter oil seed rape, 66 of beet and 57 of maize) were distributed across the UK according to where the crops are typically grown (Champion et al., 2003). Here, we used the data from the conventionally-cultivated half of the fields. Biodiversity surveys were undertaken during the FSE during 2000-20003 and they included seed rain traps to estimate the abundance of soil-surface seeds and pitfall traps to sample soil-surface active invertebrates including carabid ground beetles (Brooks et al., 2003; Heard et al., 2003). Here we used data from all the traps away from the field edge, i.e. > 2m from the field boundary. Seed rain traps were 10cm diameter pots placed 32m into the field in four locations and contents were collected every two weeks from the anthesis of the first weed species until harvest of the crop (Heard et al. 2003). The seed rain was quantified as the sum of viable seeds from all traps over this period to give the density of the total of seeds available at the soil surface during that season (*D*_*i*_). Not all seeds were identified to the species level, so for consistency we aggregated seeds at the genus level. The invertebrate pitfall traps were 6cm pots placed along four transects at 8m and 32m into the field for two-week periods and for consistency with the data from the seed rain traps we considered counts during the growing season of the crops (April/May and June/July for winter-sown crops and July/August for spring-sown crops). Carabids from the pitfall traps were identified to the species level.

For the pipeline to construct inferred food webs we needed data on the mass of seeds and carabids and the energy content of seeds. We used an existing dataset to obtain data on seed mass for all species and for seed energy content for most species (Gibbons et al., 2006). Of the 82 genera of seeds in the FSE dataset, we had information on seed energy content (kJ g^-1^) of species in 60 genera and averaged this across species within each genus. For the 22 genera without this information, we used the average energy content across genera (18.77 kJ g^-1^, interquartile range: 16.04 – 21.03 kJ g^-1^), although initial exploration indicated that using the average did not make substantial differences to the results.

Carabid body mass was derived from body length. The body length for each of the 91 carabid species was calculated as the geometric mean of the maximum and minimum length (in mm) reported by Luff (2007), and was converted to body mass (*M*_*i*_; dry mass in mg): log(*M*_*j*_) = -3.4 + (2.6 × log(body length) (Jarosik, 1989; Saska, Honěk, & Martinková, 2019).

Carabid pitfall data is a measure of activity-density, i.e. a combination of density and activity (Thomas, Brown, & Kendall, 2006), rather than true density. Activity is affected by many factors, including temperature, vegetation cover and body size (because large beetles walk further in a set time and so are more likely to encounter the pitfall trap). The relationship with body size is complex and not easy to predict (Halsall & Wratten, 1988), but recently Engel et al. (2017) used allometric scaling to predict true carabid density (*D*_*j*_) from abundance in pitfall traps (*n*_*j*_) and body mass (*M*_*j*_). The mass-specific correction factor (*β*) was related to temperature and the arrangement of the traps. Here we used the correction factor for a single pitfall trap at 21°C = -0.51 (range: -0.55 to -0.45 for 15 to 27°C) to estimate the relative density of carabids from the pitfall trap data, so 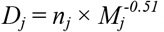. Therefore, for equal numbers of small (2mg) and large (30mg) carabids in a pitfall trap, the true density of the small carabids is four times greater than that of the large carabids.

Having constructed inferred weighted food webs for the data from 255 fields, we demonstrated the potential of this approach to be used in analysis. Firstly, we calculated the weighted connectance of the networks and tested for an effect of crop type, and including the effect of species richness and abundance of seeds and carabids as covariates. Secondly, we tested for an effect of crop type, seed size and abundance on our new network-derived metric of predation pressure, with the field as a random effect for intercept and slope to take account of variation in the size and direction of the overall effect.

## Results

### Calculation of the risk index

There was a predictable relationship of interaction cost index according to seed and carabid size. The specific result, with data points weighted by the inverse of seed size and the source of the data (Honek et al. (2003) or Petit et al. (2014)) as a random effect was:

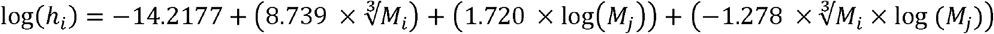

This shows that the interaction cost index for large seeds decreased with carabid size, but increased for small seeds (Fig. S1). The risk index, calculated by combining the interaction costs with the seed energy content, showed that larger beetles prefer larger seeds and have a wider diet breadth (Fig. 2), as previously found in experimental studies (Honek et al., 2003, 2007, 2007; Petit et al., 2014).

**Fig. 2.**
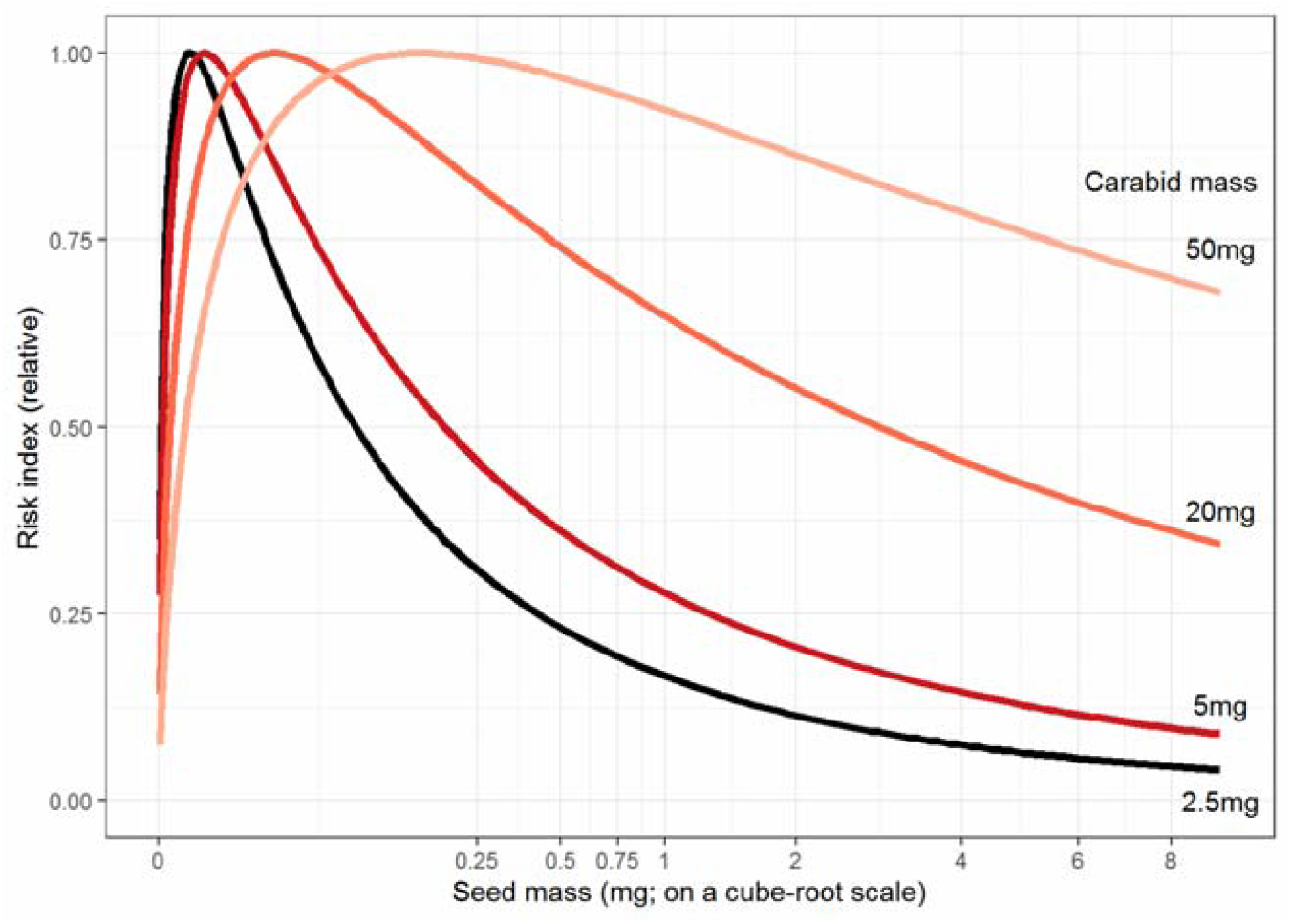
The modelled risk index (*r*_*ij*_) for seeds and four sizes of carabid (ground beetle). The risk index has been normalised so the maximum for each species is one.

### Estimation of the energetic intake of carabids

The daily energy intake reported in experimental studies varied significantly by feeding guild when seeds are provided *ad lib*. The average daily consumption rates (in mg seed per mg carabid body mass) was 0.51 for granivorous species, 0.18 for omnivorous species and (as expected) 0.00 for the single carnivorous (Fig. 3). The difference between omnivores and granivores was significant: -0.32 ±0.09 standard error in the mixed effects model, showing that omnivorous species ate fewer seeds than similarly-sized granivorous species, at least in short-term experimental studies, even when seeds were the only food available.

**Fig. 3.**
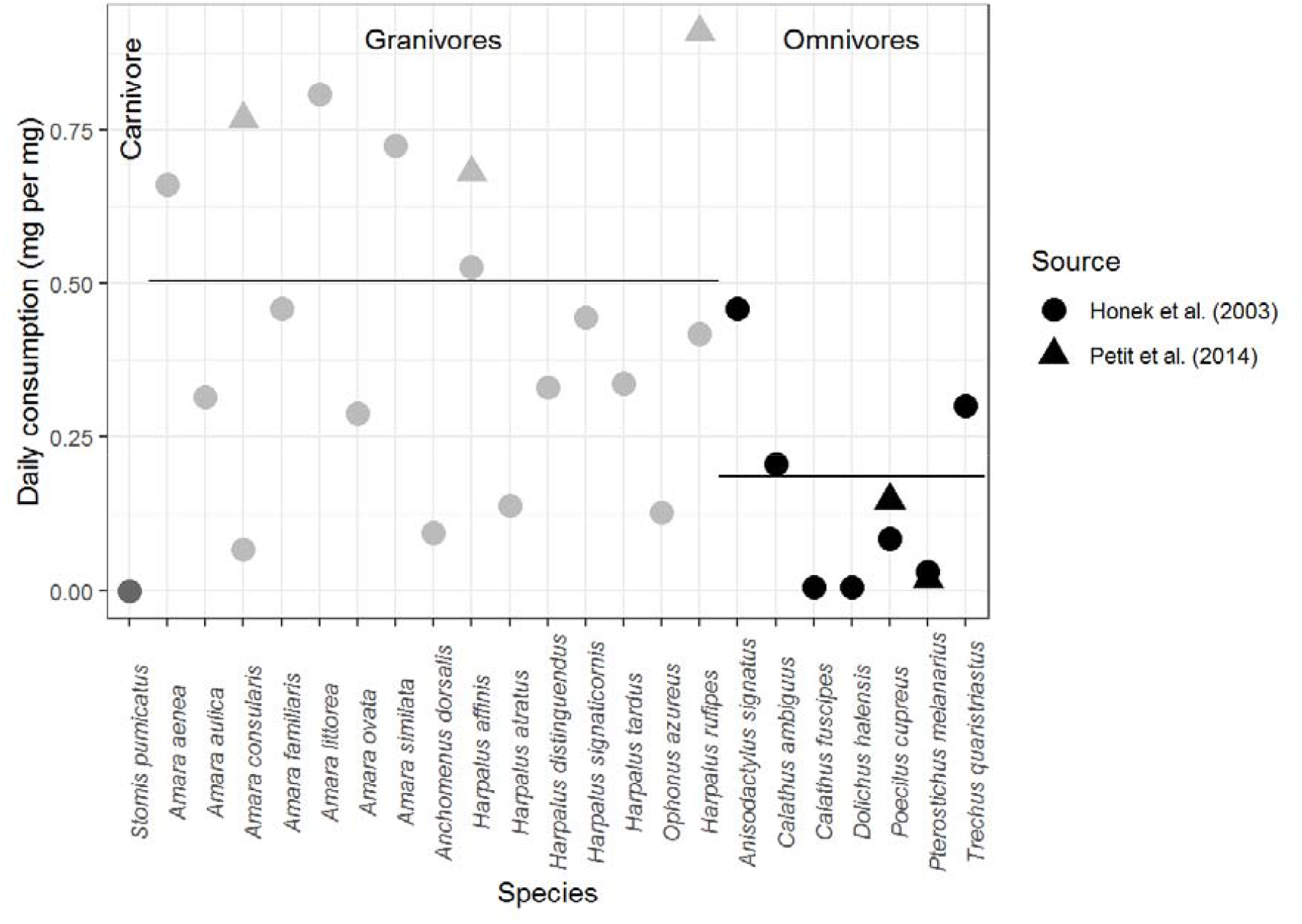
The estimates daily consumption rate of seeds by carabid beetles, from two laboratory studies, varied by their recorded feeding guilds. The horizontal lines show the mean across species in each guild, taking the source of data into account.

### Scaling up to a weighted network

We scaled up the data to construct the predicted food web for each field in this dataset, and exemplified this for a single site (Fig. 4; “WR19”: a field that growing winter-sown oilseed rape, which had the highest number of seed genera + carabid species (19 + 21) out of all sites (median = 17)). The risk index (individual seed-individual carabid) was calculated for every seed-carabid combination (Fig. 4c). There was slight variation in the risk index between similarly-sized seeds because the energy content (kJ g^-1^) varied between seeds. From the subset of seed:carabid combinations of the risk index for this site (Fig. 4d), we predicted the frequency of seeds in an individual carabid’s diet (Fig. 4e), multiplied this by the daily consumption rate (Fig 3) to give the consumption per carabid individual (Fig. 4f), and multiplied this by the relative abundance of each carabid species to give the inferred pairwise interaction strength (Fig. 4g & h).

**Fig. 4.**
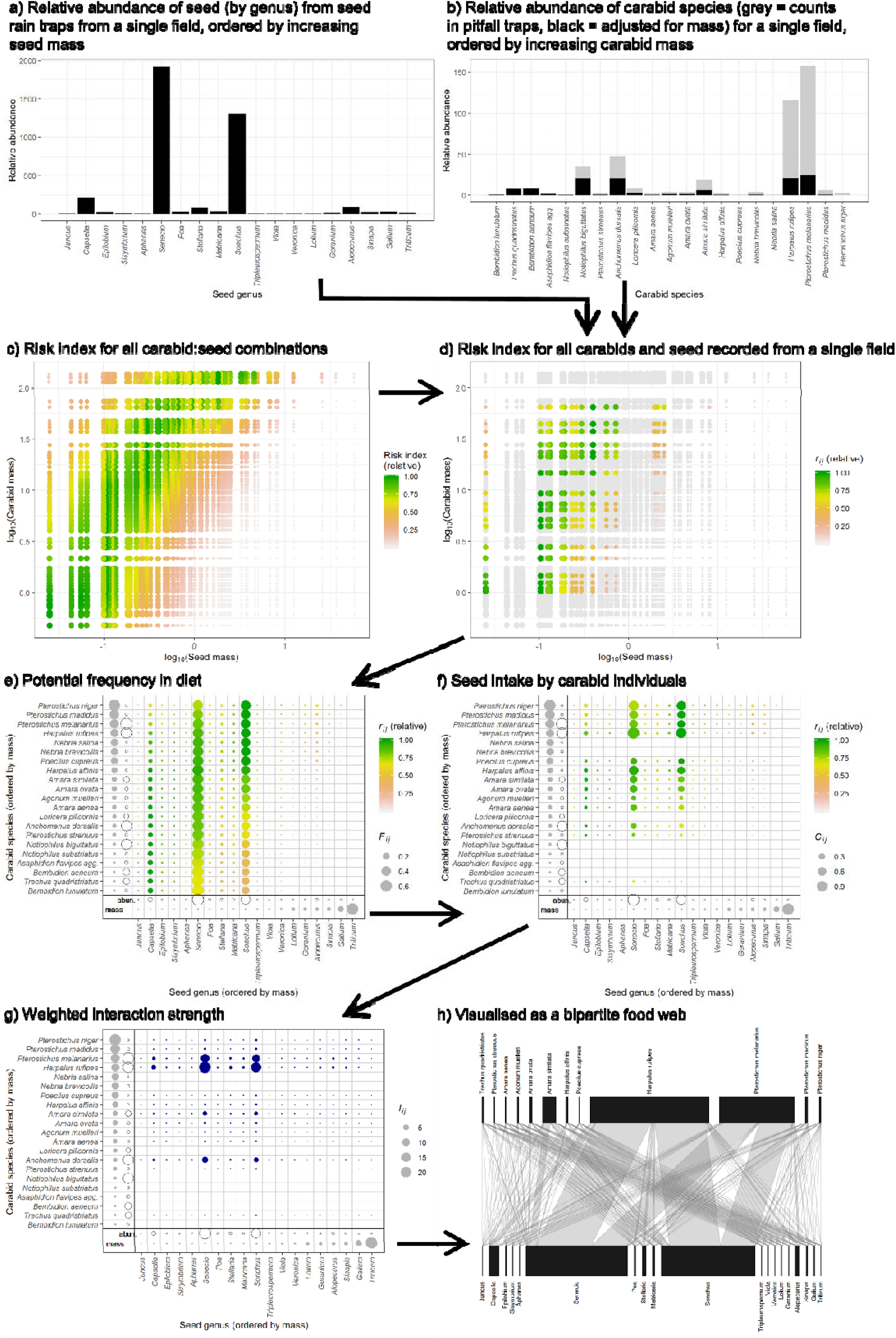
Demonstration of the pipeline for scaling up from the risk index from every seed:carabid combination to the qualitative network for a single site based on the ecological census data for carabids and seeds and the predictive model for the risk index, as derived from data in the literature. In e, f and g the abundance of the carabids and seeds is indicated by open circles and their mass is indicated in the figure margin by grey circles.

Inferred networks were constructed for all 255 fields in our dataset in which carabids and seeds were sampled: ten fields had zero carabids and eight had zero seeds, so were excluded from further analysis. The overall structure of the network, assessed with weighted connectance, was strongly affected by the network size (number of seed genera + number of carabid species; P<0.001) and abundance of seed-feeding carabids (P<0.001), but not by the field type (P = 0.394) or the total abundance of weed seeds (P = 0.255; Fig. S2). The predation pressure for each seed genus in each field varied substantially: smaller seeds tended to have lower predation pressure than larger ones (overall effect size = 0.084±0.018; P<0.001; Fig. S3), and it varied by the crop type: beet fields had the highest predation pressure ratio; maize was no different to beet (effect size ± standard error compared to beet = -0.263±0.199), spring-sown oilseed rape was lower (−0.826±0.183 compared to beet) and winter-sown oilseed rape was lowest (−1.443±0.185 compared to beet). Suitable empirical data are not currently available to validate these models, but these summary trends do provide testable hypotheses about the network-derived ecosystem function of carabids in these fields.

## Discussion

One challenge for using ecological network analysis in applied research, such as agroecology or conservation ecology, is the lack of availability of information on interactions. Here we used a mechanistic approach to infer weighted predator-prey food webs from ecological survey data and species’ traits. It was valuable to develop this method for carabid beetles because they are mobile predators of invertebrate pests and weed seeds in arable crops. In particular they are known to contribute to weed seed regulation in agro-ecosystems (Bohan, Boursault, et al., 2011; Honek et al., 2003; Kulkarni, Dosdall, & Willenborg, 2015), and network approaches have been recommended to study the direct and indirect effects of carabid biocontrol (De Heij & Willenborg, 2020). The key novelty of our approach, compared to other approaches (Morales-Castilla et al., 2015) is that we estimated interaction strength, scaled up from per capita estimates of interactions. Our approach can be used whenever the risk index for predator-prey interactions can be predicted (e.g. based on trait-matching) and where ecological census data exists, thus allowing us to create predicted food webs in places and at times when the interactions were not studied.

One key question from this research is: are these inferred networks true? It is challenging to answer this question because information on carabid-seed feeding is difficult to obtain from the field. As with any model, though, our results provide a null model that can be tested against data, when it becomes available. Our models were simple in assuming that there was bottom-up control of the carabids and assuming no interference in the field due intra-guild competition between carabids for seed resources (Carbonne, Bohan, Charalabidis, & Petit, 2019). This null model approach may show how our mechanistic models need to be refined, e.g. including the effect of traits such as seed shape, integument thickness or lipid content (Gaba, Deroulers, Bretagnolle, & Bretagnolle, 2019; Honek et al., 2007; Sebastián-González et al., 2017), or other local drivers, e.g. total seed abundance, vegetative cover or the presence of alternate invertebrate prey. Also competition for other prey or density dependence in the risk index would lead to prey switching (Gendron, 1987; Manly, Miller, & Cook, 1972). When suitable data are available, each of these assumptions could be tested in future models. Suitable data on carabid seed-feeding are likely to become widely available soon via DNA-based analysis of gut regurgitates of carabids (Sint, Guenay, Mayer, Traugott, & Wallinger, 2018; Wallinger et al., 2015) and already, simple food webs have been constructed from DNA analysis of carabid gut contents (Frei, Guenay, Bohan, Traugott, & Wallinger, 2019). Therefore when these data are combined with traditional ecological sampling of the abundance of carabids and soil-surface weed seeds, it will enable models to be validated and refined, thus providing a test of the underlying mechanisms leading to the assembly of food webs.

The impact of alternate prey would be especially important for the omnivorous carabid species that can switch to prey upon invertebrates such as slugs or aphids (Bohan et al., 2000), but even for these species, seeds are likely to be a major source of food (Frei et al., 2019). We suggest that the network model we have developed could, with care, be extended to include invertebrates as an alternate prey if the risk index could be estimated for invertebrates as well as seed prey (Roubinet et al., 2018). This would provide a powerful framework for predicting biocontrol across the community of granivorous and omnivorous carabids (De Heij & Willenborg, 2020; Ma et al., 2019).

Another test of the value of these inferred networks would be to consider the relationship between network structure and ecosystem properties (Ma et al., 2019). Here, we developed a new metric derived from the quantitative network structure: the predation pressure ratio, to explain the predicted predation rate relative to the seed abundance. The significance of this network-derived metric could be tested in the field, e.g. with bioassays of seed consumption via seed cards (Menalled, Marino, Renner, & Landis, 2000).

In our study we were able to re-use available data to model the interaction cost index (*h*, and hence risk index *r*; Fig. 3) and energetic intake per individual (*E*; Fig. 4), and so did not need to undertake our own experiments. Our reanalysis of data from the literature indicated that carabid feeding does broadly adhere to mechanistic expectations (Fig. S1) and our results matched previous qualitative expectations. If these models were extended to other organisms, there may be a lack of data in the literature, although an allometric model of handling time has previously been used to infer bird-seed feeding interaction strengths (Pocock et al., 2012). If new experiments are required to parameterise the models, it is strongly recommended that standardised experimental approaches should be used (Deroulers, Gauffre, Emeriau, Harismendy, & Bretagnolle, 2020), thus enabling results from multiple experiments to be comparable.

We note that optimal foraging theory (Pyke, Pulliam, & Charnov, 1977) is an alternative approach to Gendron’s (1987) frequency-dependent prey selection that we used to calculate the risk index, but optimal foraging models require that its equation denominators are in identical units, e.g. handling time and search time in seconds, movement in metres per second, prey density per metres squared. Although it is possible to directly assess handling time of seeds by carabids (Charalabidis, Dechaume-Moncharmont, Petit, & Bohan, 2017), such data will be available only in very well-studied systems and so we suggest that simpler models are likely to remain more appropriate for network inference in most cases.

Ultimately, the inference of interactions and the construction of potential food webs would extend the use and value of existing ecological census data. The inferred networks could then be used in ecosystem modelling or as null models to improve mechanistic understanding of species interactions. The ecological census data we used here was two decades old and, so where suitable ecological census data exist, it could be possible to construct historical inferred networks from ecological census data, and for guilds where interaction data is difficult to obtain directly, such as many insectivorous predators. The inference of weighted interactions therefore greatly extends our potential to use ecological network approaches at times and places when the interactions were not directly sampled to reconstruct the ‘interactions past’ (Bohan et al., 2017).

## Data accessibility

Code and data to replicate the figures in this paper are available at http://doi.org/10.5281/zenodo.4252783. The raw FSE data can be requested from the Environmental Information Data Centre (https://catalogue.ceh.ac.uk/documents/876358e4-62f7-4386-99e1-7d3eac223e03).

## Author contributions

MP and DB conceived the idea, MP developed the methodology, analysed the data and led the writing of the manuscript, DB and RS synthesised data, all authors critically contributed to the methodology and to drafts of the manuscript.

## Acknowledgements

MP and RS were funded by Defra (contract SCF0313). DB was funded by the Agence National de la Recherche (ANR). This work is an output of the PREAR project (Predicting and enhancing the Resilience of European Agro-ecosystems to environmental change using crop Rotations) which was a partnership between INRA, CEH, Universities of Copenhagen and Aarhus, Solagro and Szent István University. It was funded as part of the European FACCE SURPLUS (sustainable and resilient agriculture for food and non-food systems) ERA-NET co-fund scheme of Horizon 2020 programme formed in collaboration between the European Commission and a partnership of 15 countries in the frame of the Joint Programming Initiative on Agriculture, Food Security and Climate Change (FACCE-JPI). We thank David Gibbons and Jeremy Wilson (RSPB) for collating and sharing the information on seed energy content.

## Supplementary information

**Fig. S1.**
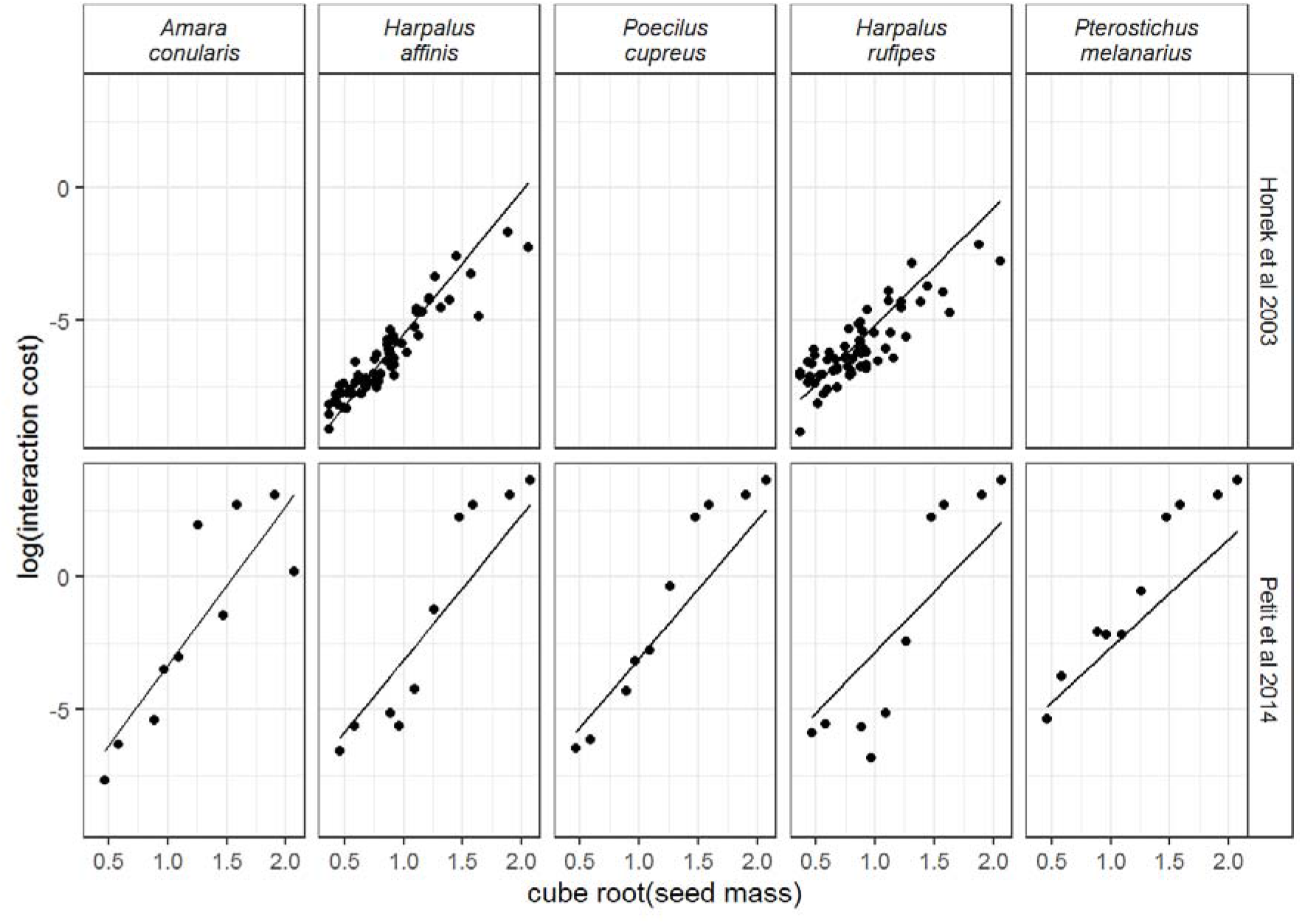
The interaction cost estimated from laboratory studies of seed choice by carabid (ground beetle) species. Carabids are arranged left to right by increasing body size (dry mass = 8.4, 13.4, 15.6, 29.6, 36.8 mg). Points are the data from published studies (Honek et al., 2003; Petit et al., 2014) and the line shows the fit of the model with seed mass, carabid mass and their interaction, with data points weighted by the inverse of cube-root of seed size.

**Fig. S2.**
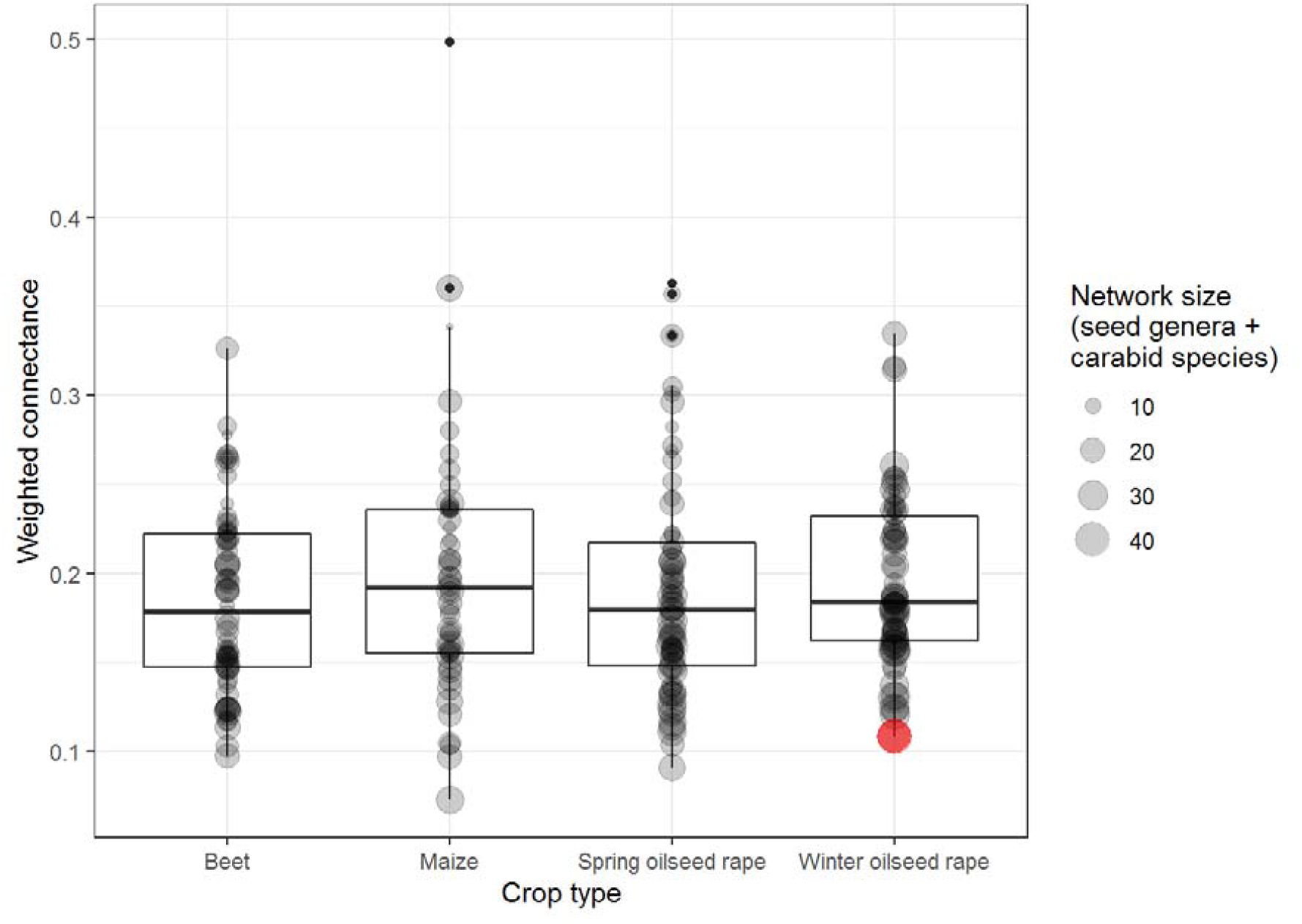
There was no association of the network structure (weighted connectance) with crop type, but it was negatively related to network size (number of seed genera + carabid species, as shown by the point size) and the abundance of seed-feeding carabids. The red point shows the data from field ‘WR19’, as shown in Fig. 4 in the main text.

**Fig. S3.**
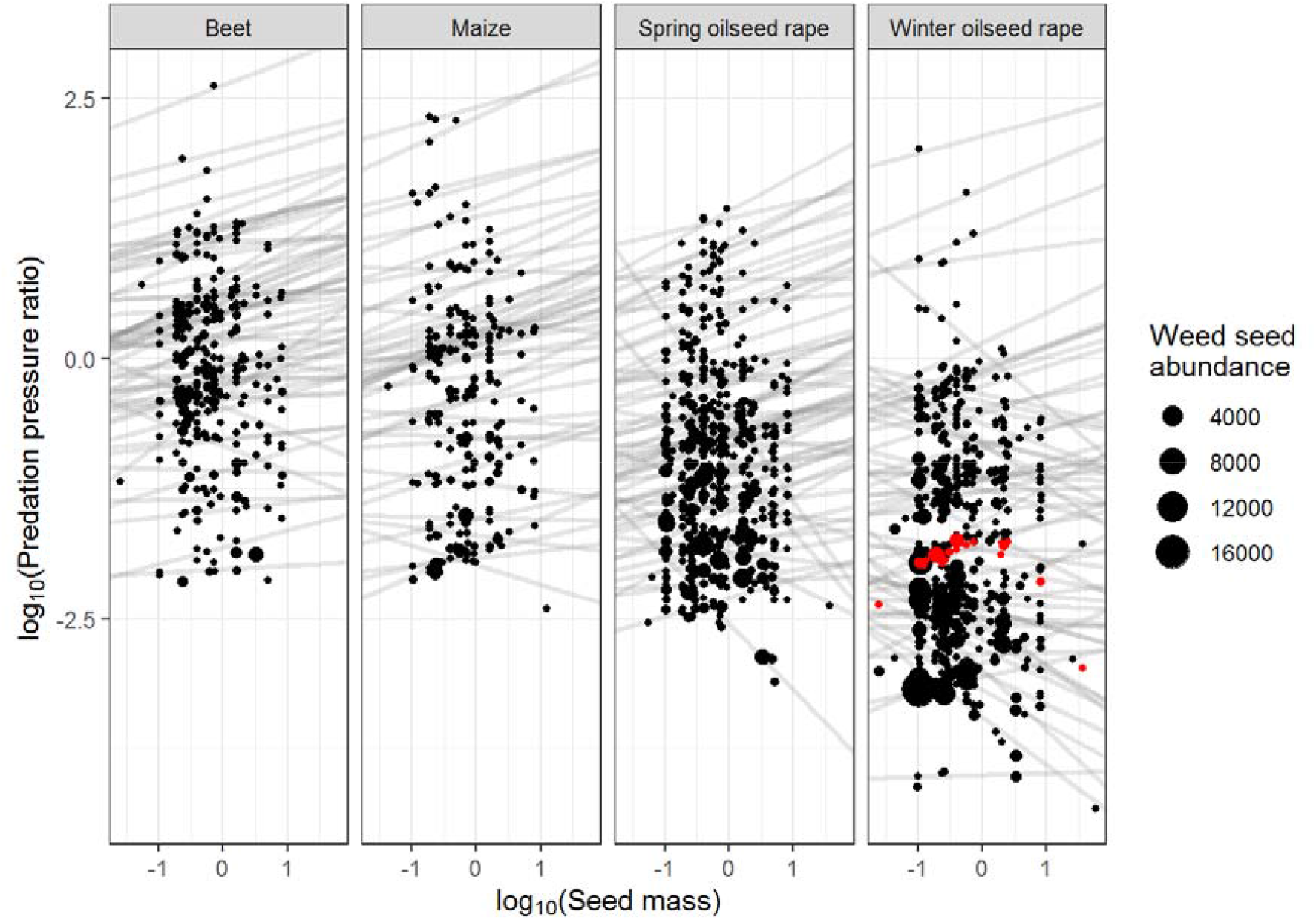
There was a clear effect of both seed mass and crop type on the predation pressure ratio. Each point represents a single seed type in a single field. Field was included as a random effect of both the intercept and the slope of the relationship. The red points show the data from field ‘WR19’, as shown in Fig. 5 in the main text.

